# Decoding of neural data using cohomological learning

**DOI:** 10.1101/222331

**Authors:** Erik Rybakken, Nils Baas, Benjamin Dunn

## Abstract

We introduce a novel data-driven approach to discover and decode features in the neural code coming from large population neural recordings with minimal assumptions, using cohomological learning. We apply our approach to neural recordings of mice moving freely in a box, where we find a circular feature. We then observe that the decoded value corresponds well to the head direction of the mouse. Thus we capture head direction cells and decode the head direction from the neural population activity without having to process the behaviour of the mouse. Interestingly, the decoded values convey more information about the neural activity than the tracked head direction does, with differences that have some spatial organization. Finally, we note that the residual population activity, after the head direction has been accounted for, retains some low-dimensional structure which is correlated with the speed of the mouse.

## Introduction

The neural decoding problem is that of characterizing the relationship between stimuli and the neural response. For example, head direction cells [1] respond with an elevated activity whenever the animal is facing a specific direction. To be able to determine this relationship, however, an experiment must be designed such that the relevant behavior of the animal can be properly sampled and tracked. It could be argued that in the case of head direction cells, the response is not too difficult to observe and characterize, but what if the stimulus driving the neural activity would have been related to specific vestibular inputs or to the head direction relative to the body of the animal. Indeed, it is not hard to imagine relevant stimuli in an open field experiment that are difficult to track and characterize, such as eye movements, limb movements, odor representations and so on. We would therefore prefer to skip the problem of devising and testing all the infinite ways of tracking and processing the behavioral data and instead allow the neural data to speak for themselves. This may seem like a daunting task but here we demonstrate that it is doable.

Our method is made possible by recent advances in recording technology that permit simultaneous recordings of a large number of neurons. While the possible states of these neurons form a very high-dimensional space, one expects that the neural activity can be described by a smaller set of parameters [2]. Dimensionality reduction techniques such as principle component analysis (PCA) and factor analysis (see [3] for a comprehensive review) can be used to obtain a lowdimensional version of the data. Here we use *persistent cohomology* and *circular parametrization* [4, 5] to identify the shape of this underlying space and then decode the time-varying position of the neural activity on it.

In summary, our method consists of combining four key steps:

1. Dimensional reduction using PCA
2. Feature identification using persistent cohomology
3. Decoding using circular parametrization
4. Removing the contribution of the decoded features on the data using a generalized linear model (GLM)

This procedure is quite general and may qualify to be called *cohomological learning* ^1^. Steps 1-3 are illustrated in Figure 1. In the last step, the process of removing the contribution of a given feature results in a new data set to which the method can then be reapplied, in a way similar to [6], in order to characterize additional features in the neural activity.

**Figure 1:**
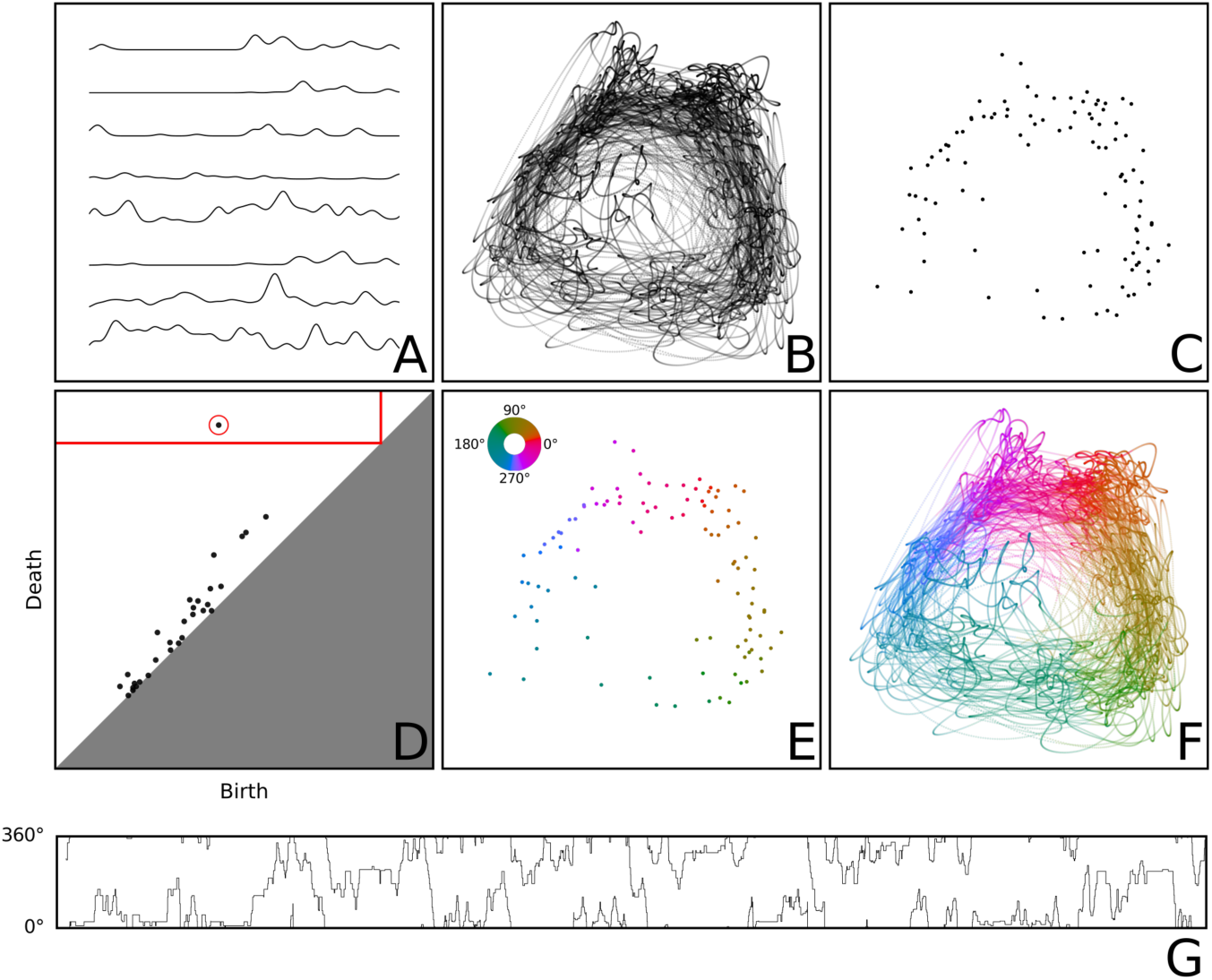
The decoding procedure. We start with an estimated firing rate for each neuron (A) obtained by smoothing the spike trains of the neurons. The firing rates are sampled at fixed time intervals, giving a point cloud (B), which is then simplified (see Appendix C), obtaining a reduced point cloud (C). The first persistence diagram (D) of the reduced point cloud is computed. We select the longest living feature (circled in red) and a scale where this feature exists (red lines). Circular parametrization is used to obtain a circular map (E) on the reduced point cloud, where the color of a point represents its circular value according to the color wheel in the upper left corner (the same coloring scheme will be used throughout the rest of the article). The map is extended to the full point cloud (F) by giving each point the value of its closest point in the reduced point cloud. In (G) we show the decoded circular value as a function of time. The point clouds in B, C, E and F are displayed as 2D projections.

As an example, we apply this method to neural data recorded from freely behaving mice [7, 8], discover a prominent circular feature and decode the time-varying position on this circle. We demonstrate that this corresponds well with the head direction, but with subtle, interesting differences. Finally, we consider the remaining features, following the removal of the head direction component, and find that there is still a structure there which is correlated to the speed of the mouse.

## Results

### Rediscovering head direction cells

We applied our decoding procedure, as summarized in Figure 1, on neural recordings of mice [8, 7]. This revealed in several cases a prominent circular feature that was then decoded as shown in Figure 2 and 8, resulting in a 1-dimensional circular time-dependent value. This decoded trajectory corresponded very well to the tracked head direction, as shown in Figure 3 and Video 1.

**Figure 2:**
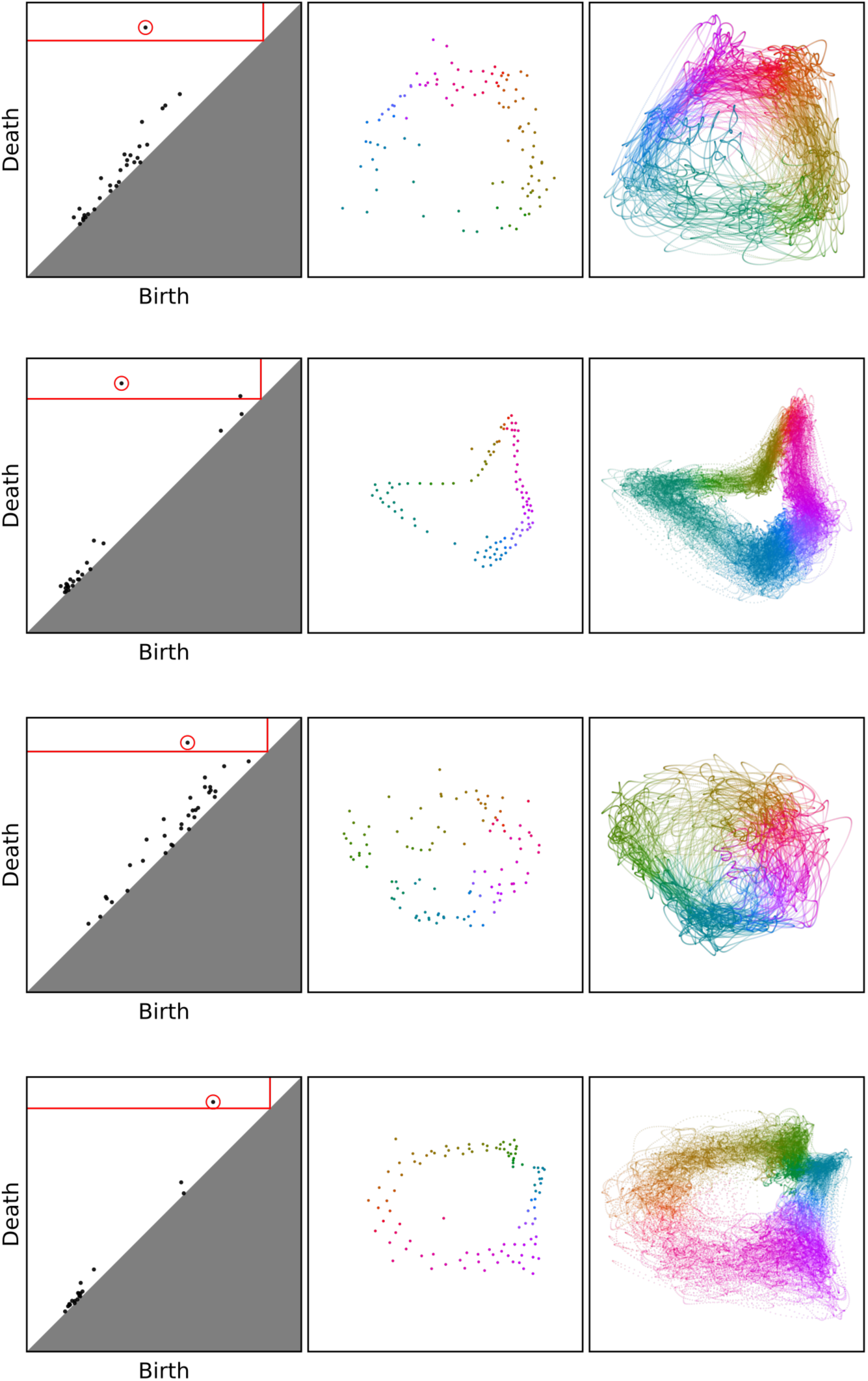
The decoding procedure applied to two recordings. From top to bottom: mouse28-140313 (using all neurons and using only identified selective neurons), mouse25-140130 (using all neurons and using only identified selective neurons). For each round we show the persistence diagram, the decoded circular value on the reduced point cloud and the circular value on the full point cloud. The point clouds in the second and third column are displayed as 2D projections.

**Figure 3:**
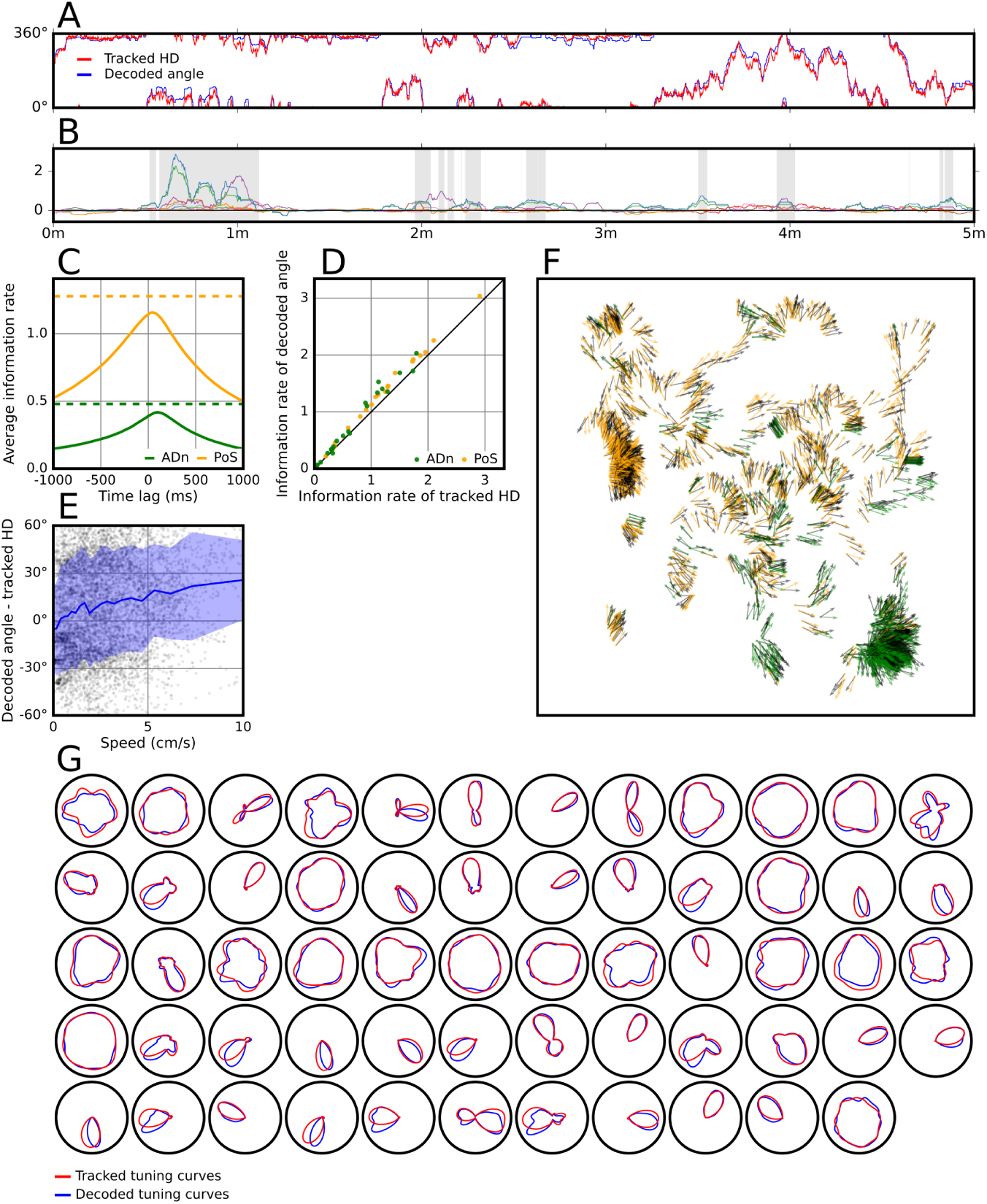
Result of the decoding procedure on mouse28-140313 (using only the identified selective neurons). A) Tracked HD and decoded direction for the first 5 minutes. B) Difference between loglikelihood of observed spike trains given decoded direction and tracked HD over individual neurons. The shaded time intervals represent the moments of drift. C) Average information rate over all ADn neurons (green lines) and PoS neurons (orange lines) of the tracked HD at different time lags (solid lines) and of the decoded direction (dashed lines) at 0 time lag. The average information rate of the tracked HD peaked at 95ms over ADn neurons and at 40ms over PoS neurons. D)Information rates of the tracked HD over all PoS neurons at 95ms time lag and over all ADn neurons at 40ms time lag, which is when their average information rates peaked, against information rates of the decoded direction at 0 time lag. E) Angular difference (decoded angle tracked HD) for each time step during drift, plotted against the speed of the mouse. We also display the average angular difference (blue line) *±*1 SD (shaded region). F) For each time step during drift, we plot the tracked HD (black arrows) and the decoded angles, shown as a green (resp. orange) arrow when deviated to the clockwise (resp. counterclockwise) of the tracked HD. The arrows are rooted at the position of the mouse at that time step. G) Tracked and decoded tuning curves of all neurons.

In detail, for each mouse we ran the procedure two times. First we ran the procedure using a large smoothing width of *σ* = 1000*ms* such that only features with slow dynamics remain, while arguably less interesting features such as local theta phase preferences are ignored. Having decoded the feature, we used an information theoretic measure to determine which cells in the population were selective for this feature [9]. We then ran the procedure again with only the spike trains of the selective neurons. This time using a smaller smoothing width of *σ* = 250*ms* to allow for a finer decoding.

In Figure 3B, C and D we see that the decoded trajectories actually convey more information about the neural activity than the tracked head direction does. We were able to resolve moments of drift during the experiment, i.e. moments when the neural data are better explained by the decoded angle than by the tracked head direction. This was done by evaluating the time-varying log-likelihood ratio of two generalized linear models (GLM) [10], one with the decoded angle and a second one with the tracked head direction. The difference of these two log-likelihoods are shown in Figure 3B for each neuron. The GLM test found 90 moments of drift, lasting 2.01 seconds on average, that were dispersed throughout the experiment. Consistent with our findings, drift of the head direction representation has been previously reported in rodents [11, 12, 13, 14].

Figure 3E shows that the discrepancy is mostly independent of the speed of the mouse, with a slight clockwise deviation at slow speeds and a slight counterclockwise deviation at higher speeds. In Figure 3F we observe what appears to be a spatial dependence of the deviation, where the decoded angle is skewed counterclockwise in some parts of the box and skewed clockwise in other parts. This suggests that the difference is not simply due to a random drift in the network representation [15], but rather that the internal representation is occationally distorted by the environment [16, 17, 18]. Another possible reason might be that the head direction is not precisely what is driving the neu ral activity but rather something similar such as the direction the body is facing or the direction that the animal is attending to. It could also be partially due to tracking error since the animal was tracked using a single camera at 30 frames per second and two points on the animal’s head.

### Capturing speed cells

Identification of a single feature from a neural recording is a first step. A single recording, however, could contain a mixture of cells responding to different features or even multiple features [19]. We therefore take an iterative approach, wherein features are identified using topological methods and then explained away by a statistical model to uncover any additional features [6]. Here we used the GLM to predict the neural activity given the decoded direction. We then subtracted the predicted neural activity from the original spike trains in an attempt to remove the contribution of the decoded direction on the activity. When we applied the decoding procedure on these residual spike trains, using a smoothing window of *σ* = 1000*ms*, we obtained a residual point cloud (shown in Figure 9A). This time we did not find any additional cohomological features, but the point cloud still indicates a non-random structure of the data, as shown in Figure 4A, suggesting that additional features might remain. The eigenvalues of the residual point cloud suggest that there is a one-dimensional feature that remains in the data. Two possible candidates are mouse speed and angular velocity, and Figure 4B and C indicate that both of these two candidates are encoded. By fitting a GLM including speed and a GLM including angular velocity to the data we see in Figure 4D that most of the neurons are more selective for speed than for angular velocity. Finally, we included both decoded HD, speed and angular velocity in the GLM and created a new residual point cloud as before (shown in Figure 9C). This time the eigenvalues (Figure 4E) are closer to random.

**Figure 4:**
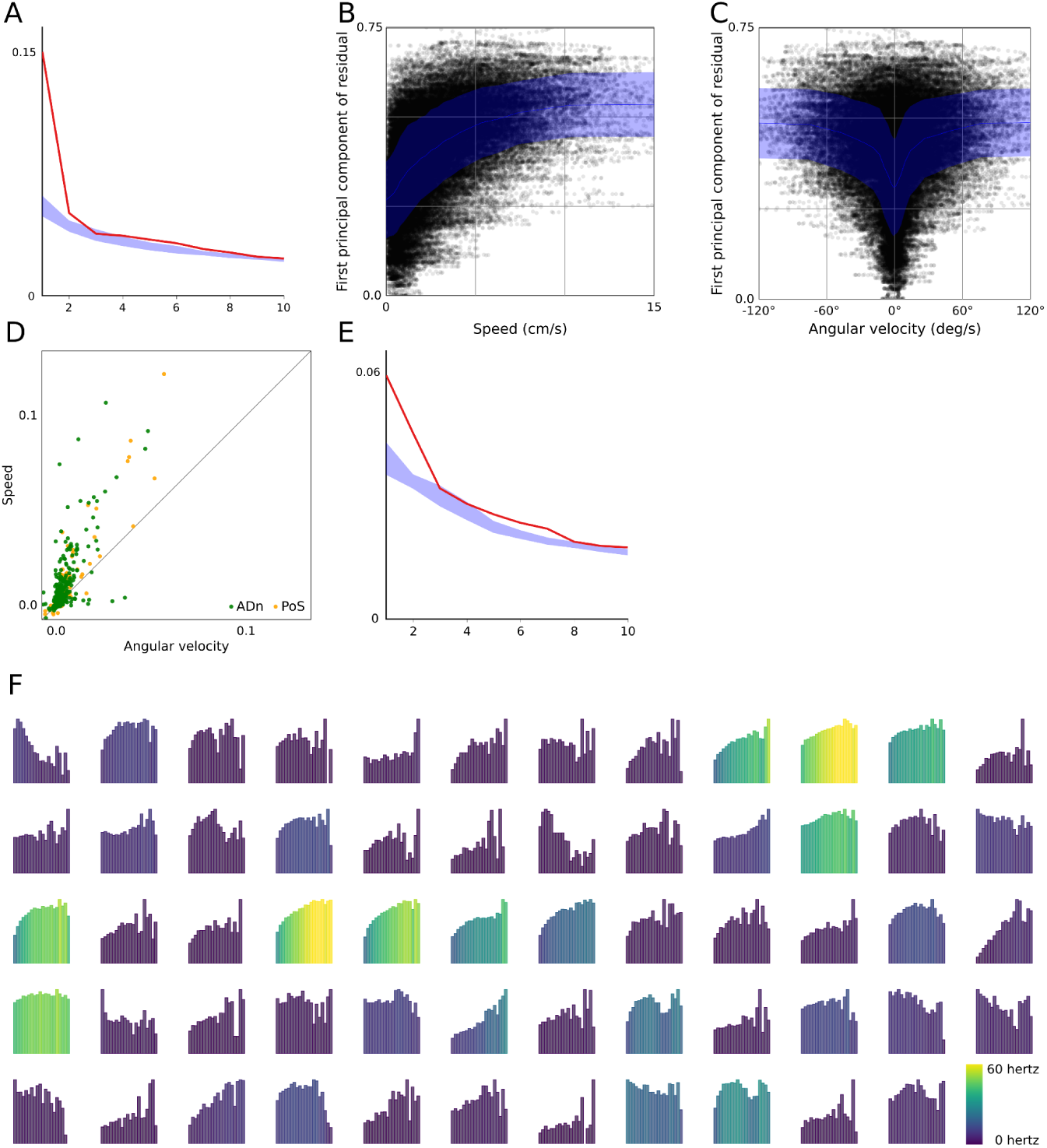
Residual analysis. A) The 10 largest eigenvalues (red line) of the residual point cloud after accounting for decoded HD, compared with a 96% confidence interval of the 10 largest eigenvalues of each of 100 shuffled point clouds (see Appendix B). B and C) The first principal component of the residual point cloud against the speed (resp. angular velocity) of the mouse, with average values (blue line) *±*1 SD (shaded region) D) The pseudo *R*^2^-score of a GLM including speed (x-axis) and a GLM including angular velocity (y-axis), averaged over a 5-fold cross-validation on all recorded neurons in the data set. E) The same as A but for the residual point cloud after accounting for decoded HD, speed and angular velocity. F) The speed tuning curves for all the neurons in mouse28140313. On the x-axis of each tuning curve is the speed of the mouse, going from 0 cm/s to 15 cm/s. On the y-axis is the average firing rate of the neuron given the speed of the mouse. Note that the y-axes are scaled indepentently for each tuning curve.

## Discussion

As illustrated in our example, the main benefit of using persistent cohomology is that it allows us to understand the shape of the neural data. It is often difficult, or even impossible, to identify the shape from a 2D or 3D projection, as seen for instance in the first round on mouse25-140130 (Figure 2), and it would likely be even more difficult for more elaborate features. We emphasize that our method does not rely on knowing that we are looking for (such as head direction). In our analysis, the recorded head direction is only used to validate the method. In practice, the discovery of a circular feature together with a circular parametrization would give the researcher useful knowledge of what kind of data could be encoded. For instance, if the circular value were instead moving in one direction on the circle with a certain frequency, this would indicate that something periodic is being encoded.

Topological data analysis has been previously ap plied to identify the shape of the underlying space in neural data [20, 21, 22]. The key contribution of our method is that we also decode the time-dependent variable on this identified space.

Some applications of topological methods to neural data [6, 22, 23, 21] have constructed spaces based on correlations between neuron pairs. This may work well when the tuning curves of the neurons are convex, but this assumption might be too strong. For instance, some of the neurons in Figure 3G are selective for two different directions. This could be due to errors in preprocesing where the activity of two neurons are combined or might even be a property of the cells themselves. Our method does not require this assumption.

Cohomological learning is not restricted to just simple features such as a circle. For instance, if the underlying space was instead a torus, this method would detect two 1-dimensional features corresponding to the two different ways to traverse a torus, as well as one 2dimensional feature. The 1-dimensional features would then give circular parametrizations that together determine the time-dependent position on the torus. Additional theoretical work, such as [24], should reveal the extent to which other features can be identified and decoded.

We were surprised to see how well the method performed with populations of less than 100 neurons and are anxious to see what insights will be gained as neural data sets of hundreds or even thousands start to be common.

## A Topological background

The state of a neural population at a given time point can be represented as a high-dimensional vector *x ∈* ℝ^*n*^. When collecting all the vectors corresponding to each time point we get what is called a *point cloud χ ⊂* ℝ^*n*^. Describing the shape of point clouds such as these, however, is non-trivial, but can be done using tools coming from the field of *algebraic topology*.

Cohomology (see Chapter 3 of [25]) is one such tool, which is able to distinguish between shapes^2^, such as a circle, sphere or torus. Intuitively, the 0-dimensional cohomology counts the number of connected components of the space, and for *n ≥* 1 the *n*-dimensional cohomology counts the number of *n*-dimensional holes in the space. For instance, a circle has one connected component and one 1-dimensional hole, a sphere has one connected component and one 2-dimensional hole, and a torus has one connected component, two 1dimensional holes and one 2-dimensional hole.

If we try to naively compute the cohomology of a finite point cloud χ *⊂*ℝ^*n*^ we run into a problem however, since this is just a finite set of points with no interesting cohomology. Instead we consider the cohomology of the *ϵ*-thickening^3^ of χ, denoted by χ _*ϵ*_, consisting of the points in ℝ^*n*^ that are closer than *ϵ* to a point in χ.

If we are able to find a suitable scale ϵ, we might recover the cohomology of the underlying shape. Often however, there is noise in the data giving rise to holes, and there is no way to separate the noise from the real features. The solution is to consider all scales rather than fixing just one scale. This is where *persistent cohomology* ^4^ [4] *enters the picture. Persistent cohomology tracks the cohomology of the space χ*_*ϵ*_ as *ϵ* grows from 0 to infinity. Such a sequence of growing spaces is called a *filtration*. When *ϵ* = 0, the set χ _*ϵ*_ is equal to χ and has no interesting cohomology. On the other hand, when *ϵ* is large enough, the space χ _*ϵ*_ becomes one big component with no holes. In between these two endpoints however, holes may appear and disappear.

The *n*’th persistent cohomology gives us the birth scale and death scale of all *n*-dimensional features. A feature with birth scale *a* and death scale *b* is denoted by [*a, b*). The features can then be drawn as a *barcode*, or alternatively, each feature [*a, b*) can be drawn as the point (*a, b*) in the plane, obtaining the *n’th persistence diagram*, as shown in Figure 5. The features far away from the diagonal persist longer in the filtration, and are considered more robust, while features near the diagonal are considered as noise.

**Figure 5:**
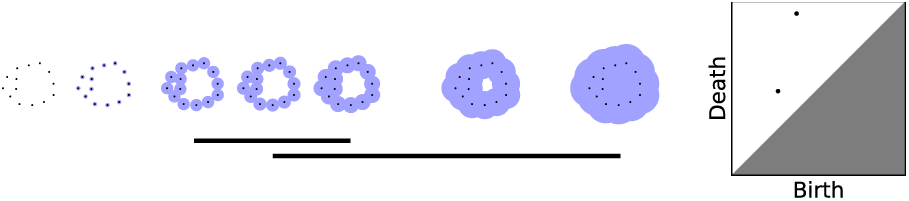
Left: A filtration together with its first persistent cohomology, drawn as a barcode. Two 1-dimensional holes appear and disappear during the filtration, as reflected in the bars. Right: First persistence diagram of the filtration.

Once the persistence diagram has been computed, we want to relate the discovered features to the point cloud. This can be done using *circular parametrization* ([4, 5], *see also Appendix D), which turns any chosen 1-dimensional feature existing at a scale ϵ* into a circle-valued function *f*: *X → S*^1^, revealing the feature. See Figure 1.

## B Details about the methods

### Decoding procedure

We will here give the details of the decoding procedure summarized in Figure 1. The parameters used in our analysis were chosen experimentally.

We are given a set 𝒩 = {1, 2, *…, N*} of neurons and a spike train 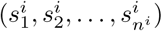 for each neuron *i ∈ 𝒩*, consisting of the time points where the neuron fires. The neurons having a mean firing rate lower than 0.05 spikes per second are discarded. The remaining spike trains are smoothed with a gaussian kernel of standard deviation *σ*, and then normalized to take values between 0 and 1. This gives us estimated normalized firing rates *f*_*i*_: ℝ *→* [0, 1] for each neuron *i*, as shown in Figure 1A. The firing rates are then sampled at fixed time intervals of length *δ* = 25.6*ms*^5^, *resulting in a sequence (****x***_*0*_, **x**_1_, *…*) of points in ℝ^*N*^, where **x**_*t*_ = (*f*_1_(*δt*), *f*_2_(*δt*), *…, f*_*N*_ (*δt*)). In order to reduce noise, we project this point cloud onto its first *d* = 6 principal components, resulting in a ordered point cloud *X ⊂* ℝ^*d*^ as shown in Figure 1B (in 2D). We then simplify (see Appendix C) the point cloud *X*, obtaining the point cloud *S*, shown in Figure 1C. The first persistent cohomology of *S* is then computed. We focus our attention on the longest living feature [*a, b*), and pick a scale ϵ *∈* [*a, b*) where this feature exists. We used ϵ = *a* + 0.9(*b - a*) in our analysis.

Figure 1D shows the persistence diagram. The chosen feature [*a, b*) is circled in red, and the scale ϵ is marked with red lines. We apply circular parametrization to obtain a circular value, i.e. an *angle θ*(*x*) for each point *x ∈ S*, as shown in Figure 1E. We then extend the function *θ* to χ by giving a point *x ∈ χ* the circular value of its closest point in *S*. The final decoding is shown in Figure 1F and G.

### Identifying selective neurons

We compute the information rate, as described in [9], for each neuron relative to the circular value using the formula

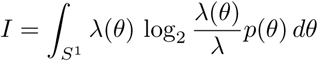

where *λ*(*θ*) is the mean firing rate of the neuron when the circular value is *θ, λ* is the overall mean firing rate of the neuron, and *p*(*θ*) is the fraction of timepoints where the circular value is *θ*. The integral is estimated by partitioning the circle in 20 bins of width 18 degrees, and assuming that the mean firing rate *λ*(*θ*) is constant in each bin. We identify a neuron as being selective if its information rate is larger than 0.2. The method can then be run again without the spike trains of the non-selective neurons. The information rates of each neuron relative to the tracked HD, shown in Figure 3C and D, were calculated in the same way.

### Comparison with recorded HD

For the comparisons with the tracked head direction in Figure 3, we rotated the decoded circular value to minimize the mean squared deviation from the tracked HD. The tuning curves in Figure 3G were made by taking the histogram of the angles at the spikes of each neuron, divide by the time spent in each angle, and smooth with a gaussian kernel of 10 degree standard deviation.

### Calculating mouse speed and angular velocity

The speed of the mouse was calculated by first smoothing the recorded position of the mouse separately in the x-axis and y-axis with a gaussian kernel of 0.1s, and then taking the central difference derivative of the smoothed position at each time step, letting *h* = 25.6*ms*. Similarly, the angular velocity was calculated by first smoothing the recorded head direction of the mouse with a gaussian kernel of 0.1s, and then taking the central difference derivative at each time step, letting *h* = 25.6*ms*.

### GLM

We fit a GLM to the activity of each neuron *i∈ 𝒩* as follows. First we binned the spike train of the neuron into bins of size 25.6*ms*, and the circle into angular bins of width 36 degrees. Let *X*_*j*_(*t*) be the indicator function which is 1 if the decoded value is in the *j*’th angular bin at time step *t* and 0 otherwise, and let *y*_*t*_ be the observed spike count of the neuron in time bin *t*. Assuming that the firing of the neuron is an inhomogeneous poisson process with instantaneous intensity at time step *t* given by

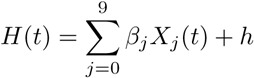

where *h* and the *β*_*j*_ are parameters, the log-likelihood of observing the spike train of neuron *i* is given by

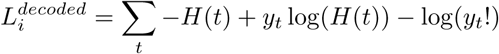

We used [30] to obtain the parameters (*h*,{*β*_*j*_}) that maximizes the log-likelihood. The log-likelihood of the neuron having *y*_*t*_ spikes in bin *t* is now given by

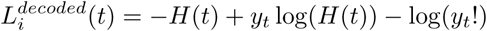

We then repeated this process, replacing the function *X*_*j*_(*t*) by the indicator function corresponding to the tracked HD, and obtained the corresponding values 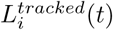. The difference 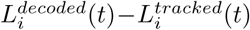 represents how much better the neuronal activity at time step *t* is explained by the decoded value compared to the tracked HD. This difference is shown for each neuron in Figure 3B. The time intervals when the sum of these differences were above a certain threshold (we used *thresh* = 1) is referred to as *moments of drift*.

### Residual analysis

Given the above GLM, we made a residual spike train for each neuron *i ∈ N* by taking the original spike train of the neuron and adding for each time step *t* a negative spike of magnitude *H*(*t*) at the center of bin *t*. This is similar to the residual spike trains described in [6].

Figure 4A was made as follows. The red line shows the 10 largest eigenvalues of the covariance matrix of the residual point cloud after accounting for the decoded direction in the GLM. Each of the residual spike trains that gave rise to this point cloud were then shifted in time by *n* time steps, where *n* is a number sampled uniformly in the range from 0 to 100. This was done 100 times, resulting in 100 point clouds. For each of the these point clouds, the 10 largest eigenvalues of their covariance matrix were chosen. The blue band in Figure 4B shows for each index *i* a 96% confidence interval of the *i*-th largest eigenvalue, sampled from this empirical distribution.

Figure 4D and 9B were made as follows. We binned the mouse speed in *n-*1 equally sized bins in the range [0, 15) and one bin [15, *∞*), and the angular velocity in *n-*2 equally sized bins in the range 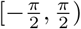 and the two bins 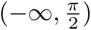 and 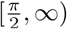. We then defined two GLMs as before letting the functions *X*_*j*_(*t*) be the indicator functions on the speed bins (resp. the angular velocity bins). These two models were cross-validated using 5 folds on each recorded neuron of every mouse in the data set, for different choises of the number of bins *n*. The average pseudo *R*^2^-score over all folds and over all neurons is shown in Figure 9B for the two models and for different values of *n*. The average pseudo *R*^2^score over all folds for each indivual neuron is shown in Figure 4D for the two models, using 20 bins in each model, which is where the average pseudo *R*^2^-scores peaked.

Figure 4E was made the same way as Figure 4A, but for the residual point cloud after accounting for decoded direction, speed and angular velocity in the GLM.

## C Point cloud simplification

The procedure of simplifying the point cloud χ to obtain the point cloud *S* consists of two steps; *Radial distance* and *Topological denoising*.

### Radial distance

To make the computations faster, we simplify χ using the following method, called *Radial distance* [31]. Start with the first point in χ and mark it as a key point. All consecutive points that have a distance less than a predetermined distance E to the key point are removed. The first point that have a distance greater than *ϵ* to the key point is marked as the new key point. The process repeates itself from this new key point, and continues until it reaches the end of the point cloud. This procedure results in a smaller point cloud χ *′* that is close to the original point cloud^6^. The parameter *ϵ* is typically determined proportionally to the spread of the point cloud. In our analysis, we used *ϵ* = 0.02.

### Topological denoising

Since the point cloud is noisy, we need to reduce the amount of noise before we can look for topological features. We use a method for topological denoising that was introduced in [32]. Given a subset *S* of χ *′*, we define the function

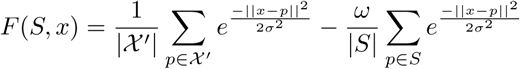

The parameter *σ* is an estimate on the standard deviation of the noise, and the parameter *w* determines how much the points in *S* should repell each other. We maximize the function *F* by iteratively moving each point in *S* in the direction of the gradient of *F*: Starting with a sample *S*_0_ *⊂χ ′*, we let

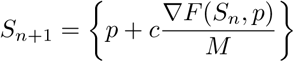

for all *n*, where

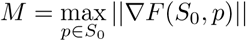

and *c* is a parameter determing the maximum distance the points in *S*_*n*_ can move. In our case, we constructed *S*_0_ by taking every *k*’th point of χ′, where *k* is the largest number such that *S*_0_ had 100 points. We used the parameters *σ* = 0.1*s, w* = 0.1 and *c* = 0.05, where *s* is the standard deviation of *X′*. We did 60 iterations resulting in the point cloud *S* = *S*_60_.

## D Circular parametrization

The material in this appendix is mostly a reformulation of parts of [4] with exception of the section *Improved smoothing*, which is our own contribution.

### Simplicial complexes

In topological data analysis, shapes are modeled by *simplicial complexes*. A simplicial complex can be thought of as a space that is constructed by gluing together basic building blocks called simplices.

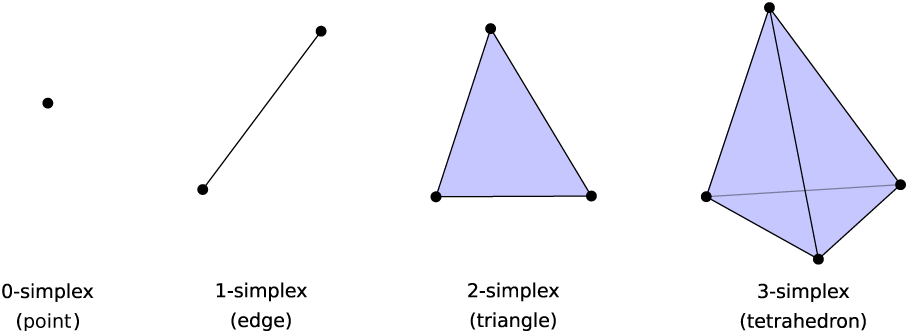

#### Definition 1.

*A* simplicial complex *is a pair* (*X, S*), *where X is a finite set called the* underlying set *and S is a family of subsets of X, called* simplices, *such that*

1. *For every simplex σ ∈ S, all of its subsets σ′ ⊆ σ are also simplices.*
2. *For every element x ∈ X, the one element set {x} is a simplex.*

An *n-simplex* is a simplex consisting of *n* + 1 elements. The 0-simplices, 1-simplices and 2-simplices are called respectively points, edges and triangles. The simplicial complexes used in our method are called *Rips complexes*, which are defined as follows.

#### Definition 2.

*Let χ⊂* ℝ^*n*^ *be a point cloud. The* Rips complex *of χ at scale ϵ, denoted Rϵ* (χ) *is the simplicial complex defined as follows:*

- *The underlying set isχ.*
- *A subset σ ⊂χ is a simplex iff ||x - y|| ≤ ϵ for all x, y ∈ σ.*

### Cohomology

Let *X* be a simplicial complex, and let *X*_0_, *X*_1_ and *X*_2_ be respectively the points, edges, and triangles of *X*.

We will assume a total ordering on the points in *X*. If {*a, b*} is an edge with *a < b*, we will write it as *ab*. Similarly, if {*a, b, c*} is a triangle with *a < b < c*, we will write it as *abc*.

Given a commutative ring 𝔸, for instance ℤ, ℝ or 𝔽_*p*_, we define 0-chains, 1-chains and 2-chains with coefficients in 𝔸 as follows:

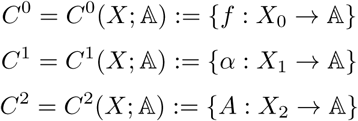

These are modules over 𝔸. We define the coboundary maps *d*_0_: *C*^0^ *→ C*^1^ and *d*_1_: *C*^1^ *→ C*^2^ as follows:

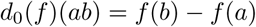

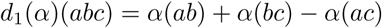

A 1-chain *α* is called a *cocycle* if *d*_1_(*α*) = 0, and it is called a *coboundary* if there exists a 0-chain *f ∈ C*^0^ such that *α* = *d*_0_(*f*). It is easy to show that all coboundaries are cocycles. We define the first cohomology of *X* with coefficients in 𝔸 to be the module

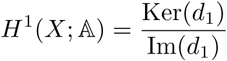

Two 1-chains *α* and *β* are said to be *cohomologous*, or belonging to the same *cohomology class* if *α − β* is a coboundary.

### Circular parametrization

The idea behind circular parametrization comes from the following theorem.

#### Theorem 1

(A special case of Theorem 4.57 in [25]). *There is an isomorphism*

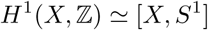

*between the 1-dimensional cohomology classes with integer coefficients of a space X and the set of homotopy classes of maps from X to the circle S*^1^ = ℝ*/*ℤ.

Given a representative cocycle *α∈ C*^1^(*X*; ℤ), the associated map *θ*: *X →* ℝ*/*ℤ is given by sending all the points in *X* to 0, sending each edge *ab* around the circle with winding number *α*(*ab*), and extending linearly to the rest of *X*. This extention is well-defined since *d*_1_(*α*) = 0.

The circular maps obtained this way are not very smooth, since all points in *X* are sent to the same point on the circle. In order to allow for smoother maps, we consider cohomology with real coefficients. Consider *α* as a real cocycle, and suppose we have another cocycle *β ∈ C*^1^(*X*; ℝ) cohomologous to *α*. Since it is cohomologous, it can be written as *β* = *α* + *d*_0_(*f*) for some *f ∈ C*^0^(*X*; ℝ). We define the map *θ*: *X* → ℝ*/*ℤ by sending a point *a* to *θ*(*a*) = *f* (*a*) (mod ℤ), and an edge *ab* to the interval of length *β*(*ab*) starting at *θ*(*a*) and ending at *θ*(*b*). This map is extended linearly as before to the higher simplices of *X*.

Given a cocycle *α ∈ C*^0^(*X*; ℤ), we now want to find a real 0-chain *f ∈ C*^0^(*X*; ℝ) such that the cocycle *β* = *d*_0_(*f*) + *α* is smooth, meaning that the edge lengths *||β*(*ab*)*||* are small. Define

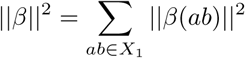

We want to minimize this value. The desired 0-chain can be expressed as

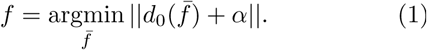

This is a least squares problem and can be solved using for instance [33].

### Improved smoothing

When we tried to apply circular parametrization to the data sets, we quickly discovered that it often produces unsatisfactory results. We demonstrate this using a constructed example, namely the Rips complex shown in Figure 6a. The first cohomology of this complex with integer coefficient is ℤ, generated by the cocycle which has value 1 on the rightmost edge, and zero on all the other edges. The color of a point indicate its circular value after applying circular parametrization.

**Figure 6:**
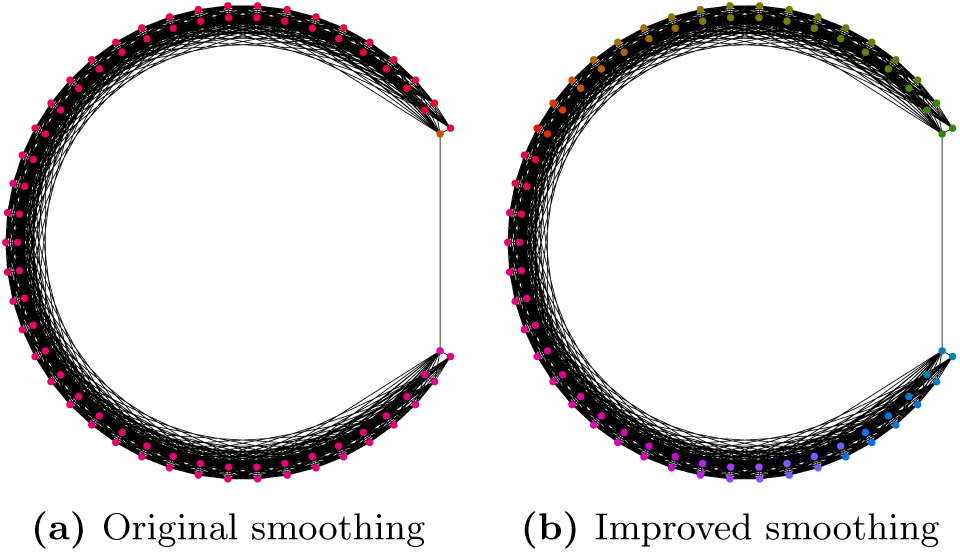
Comparison between original and improved smoothing for the constructed example.

We see that the original smoothing sends the points to circular values that are close to each other. The reason for this behavior is that the edges in the simplicial complex are not evenly distributed around the circle; there is only one edge covering the rightmost part of the circle. Since this edge gives a relatively small contribution to the sum of squares of edge lengths, it will be streched around the circle so that the other edges can get smaller. This is problematic, because we want the geometry of the complex to be preserved.

We will describe a heuristical approach to improve the situation. First, we assume that the underlying set of the simplicial complex comes with a metric *d*: *X × X →* ℝ. When the simplicial complex is a Rips complex of a point cloud χ *∈* ℝ^*n*^, we will take the metric to be the euclidean metric. Let *α ∈ C*^1^(*X*, ℤ) be the cocycle that we want to use for circular parametrizaion. Now, instead of treating every edge equally, we will assign a positive weigth *w*: *X*_1_ *→* (0, *∞*) to each edge in the simplicial complex. We define the weighting by the following procedure. First, we solve the original optimization problem (1) to obtain a cocycle *β* = *d*_0_(*f*)+*α*. We then construct a weighted directed graph *G* as follows: The vertices of *G* are the points *a ∈ X*^0^. For each edge *ab ∈ X*^1^ with *β*(*ab*) *≥* 0 there is an edge in *G* from *a* to *b* denoted *ab*, and for each edge *ab ∈ X*^1^ with *β*(*ab*) *<* 0 there is an edge in *G* from *b* to *a* denoted *ba*. This gives us a bijection between edges in *X* and edges in *G*. The weight of an edge *ab* is given by *d*(*a, b*). It can be shown that every edge *ab* in *G* has at least one directed cycle going through it. Now, for every edge in *G*, take the shortest directed cycle in the graph going through it. This gives us a collection of cycles in *G*. We now define

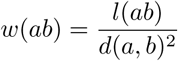

where *l*(*ab*) is the number of cycles going through the corresponding edge in *G*.

Given this weighting, we then define

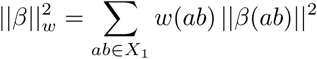

Our new optimization problem then becomes

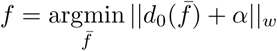

which is again a least squares problem. After solving this system, we obtain a new circular parametrization. The result of applying this procedure to the Rips complex in our example is shown in Figure 6b. Here the rightmost edge is given a much larger weighting than the other edges, since every directed cycle has to go through this edge. The result is that the circular values of the points are more evenly distributed around the circle. In Figure 7 we compare the original and improved smoothing on the circular parametrization done in the first round of mouse28-140313.

**Figure 7:**
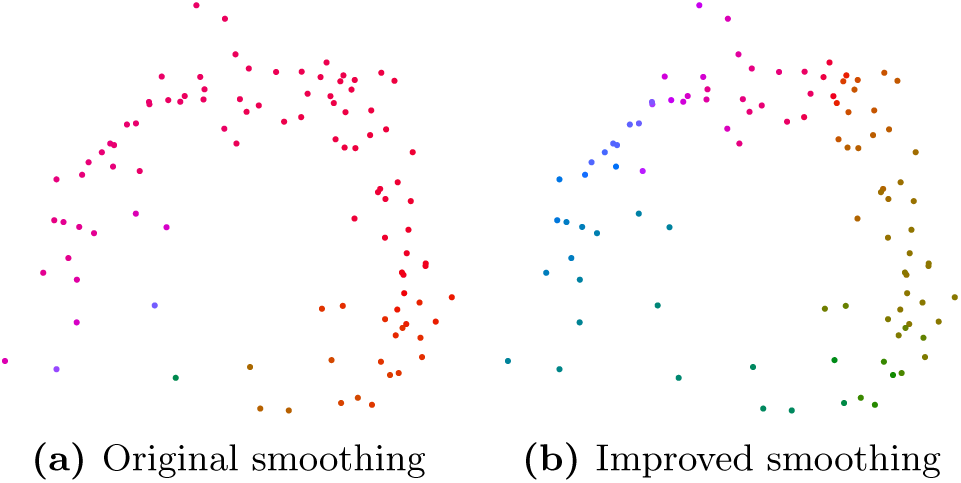
Comparison between original and improved smoothing for real data.

**Figure 8:**
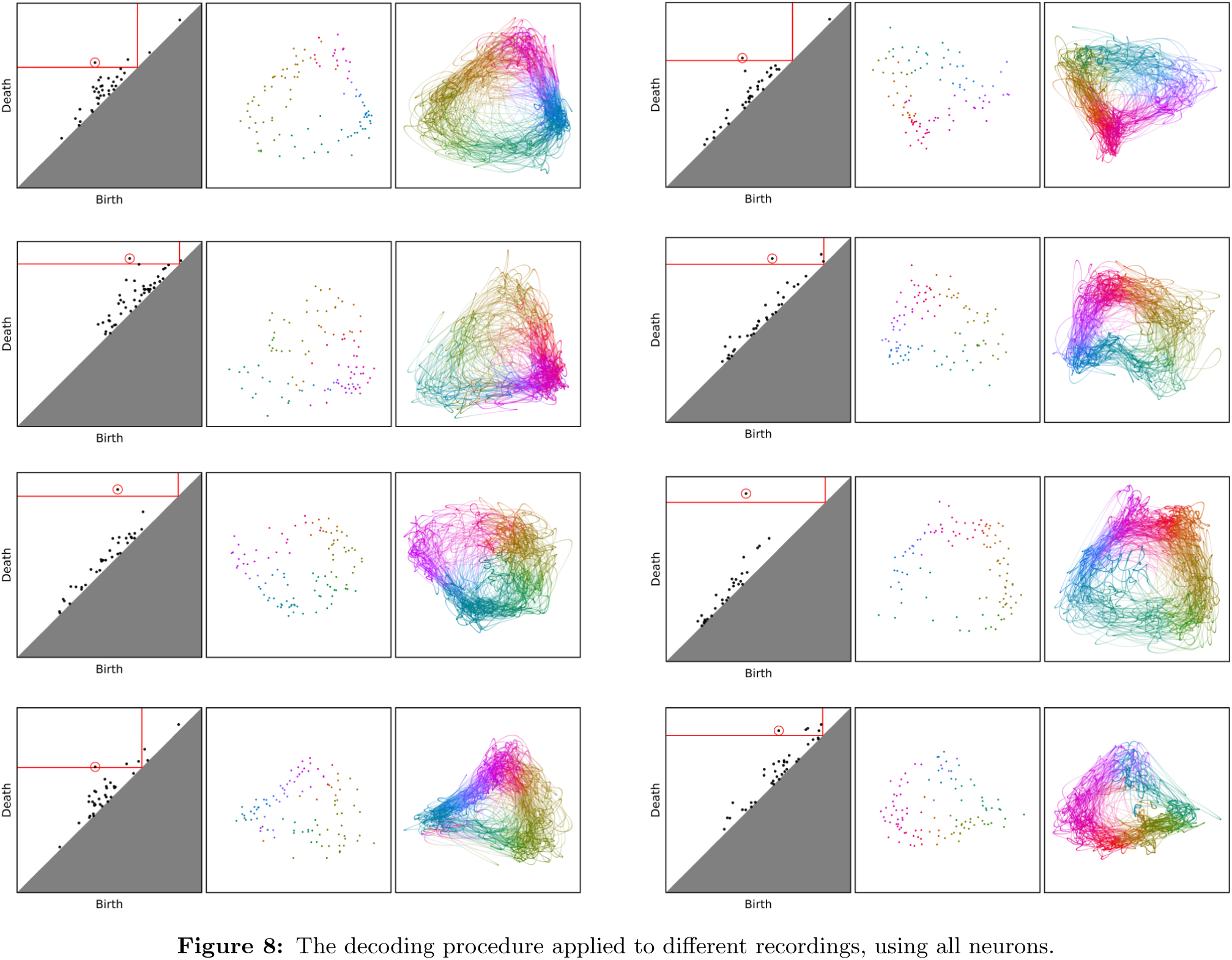
The decoding procedure applied to different recordings, using all neurons.

**Figure 9:**
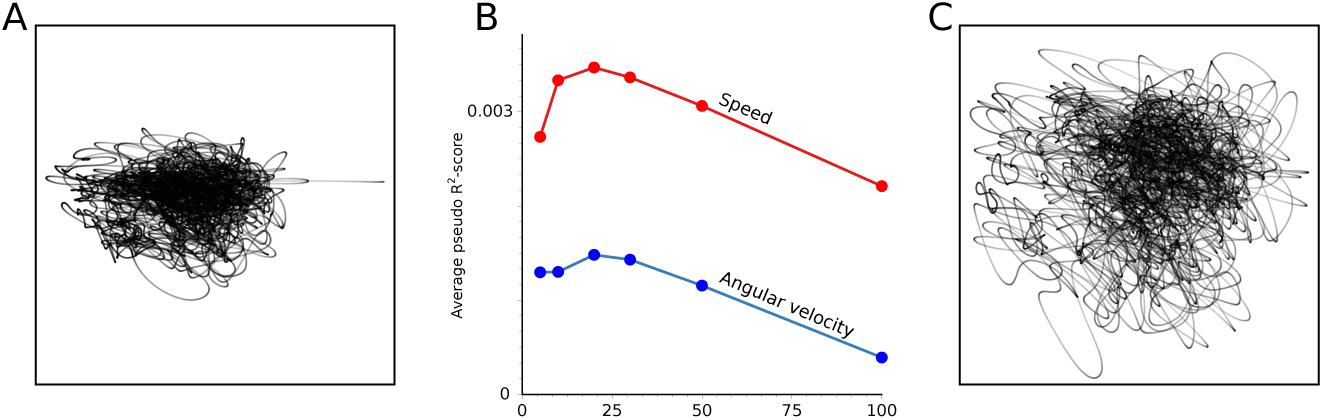
Supplementary figures to residual analysis in Figure 4. A) Residual point cloud after accounting for the decoded direction. B) The pseudo *R*^2^score of a GLM including speed (red line) and a GLM including angular velocity (blue line) for different number of bins, averaged over a 5-fold cross-validation on all recorded neurons in the data set. C) Residual point cloud after accounting for decoded direction, speed and angular velocity.

## E Persistent cohomology

By varying the parameter *ϵ* of the Rips complex of a point cloud *X*, we get a parametrized family of simplicial complexes *R*_*ϵ*_(χ), where *ϵ* goes from 0 to infinity. For every pair of parameters 0 *≤ a ≤ b* we have a natural inclusion map 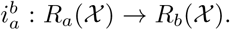. We have the following two properties:

1. The map 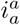 is the identity map on *R*_*a*_(χ) for all *a ∈* ℝ.
2. For all parameters *a ≤ b ≤ c* we have 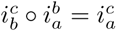.

Such a sequence of simplicial complexes together with maps satisfying the above two conditions is called a *filtration*. Let *p* be a prime, and let 𝔽_*p*_ be the field of order *p*. When taking the cohomology of each space *R*_*ϵ*_(*X*) with coefficients in 𝔽_*p*_ we get a parametrized family of vector spaces *H*^*n*^(*R*_*ϵ*_(χ)) together with the induced linear maps 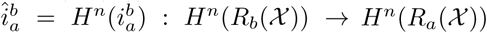 for every pair of parameters 0 ≤ *a* ≤ *b*. We have the following two properties:

1. The map 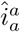 is the identity map on *H*^*n*^(*R* (χ)) for all *a ∈* ℝ.
2. For all parameters *a ≤ b ≤ c* we have 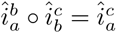.

Such a sequence of vector spaces together with linear maps satisfying the above two conditions is called a *persistence module*. The persistence module obtained by taking the cohomology of a filtration is called the *persistent cohomology* of the filtration. Another kind of persistence modules are *interval modules*.

### Definition 3.

*Let J be an interval on the real line. A J-interval module, written* II^*J*^ *is the persistence module with vector spaces*

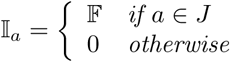

and linear maps

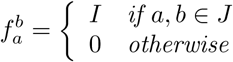

*where I denotes the identity map on* 𝔽.

### Theorem 2

(A special case of Theorem 2.7 in [34]). *Given a finite point cloud χ, the nth persistent cohomology of the Rips complex of χ can be decomposed into a sum of interval modules*

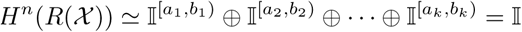

The *persistent cohomology algorithm*, described in [4], finds the intervals in the decomposition, along with a representative cocycle *α*_*i,ϵ*_ *∈ C*^1^(*R*_*ϵ*_(χ)) for every interval [*a*_*i*_, *b*_*i*_) and *a*_*i*_ *≤ ϵ < b*_*i*_, such that

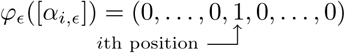

By drawing each interval [*a, b*) as a point (*a, b*) in the plane we obtain the *nth persistence diagram* of the filtration. We may now apply circular parametrization as follows. Pick an interval [*a*_*i*_, *b*_*i*_) in the decomposition. Then choose a scale *a*_*i*_ *≤ ϵ < b*_*i*_, and take the corresponding cocycle *α*_*i,ϵ*_. Now, this cocycle has coefficients in 𝔽_*p*_, but circular parametrization requires a cocycle with integer coefficients. We need to *lift α*_*i,ϵ*_ to an integer cocycle. We do this by first constructing an integer 1-chain by replacing each coefficient in *α*_*i,ϵ*_ with the integer in the same congruence class lying in the range 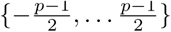. This almost always^7^ gives an integer cocycle 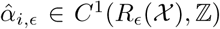. We can then apply circular parametrization to this cocycle.

In our computations, we used Ripser [35] to compute the first persistent cohomology with coefficients in 𝔽_47_, giving persistence diagrams and the associated representative cocycles. The procedure described above to lift this cocycle always gave integer cocycles.

## F Caption for Video 1 (attached)

### Decoding on mouse28-140313

We show the the position of the two LED lights that were attached to the mouse (green and orange dots), the point in between the two lights (black dot), the tracked head direction (orange line) and the decoded direction (blue line) where we decoded using only the identified selective neurons.

## Acknowledgement

We would like to thank Adrien Peyrache and Gyorgy Buzsaki for sharing the data sets [8, 7] used in our analysis for free online.

## Footnotes

1 See Appendix B for additional details.

2 Technically speaking the homotopy types.

3 In practice, we replace the ϵ-thickenings by approximations called *Rips complexes*, see Appendix D.

4 In topological data analysis, persistent *homology* is often used instead, but we will need cohomology for the decoding step. We refer to [26, 27] for introductions to persistence, and [28, 29] for introductions to applied topology in general.

5 The same interval length as were used for the camera tracking.

6 Since each point in *XI* has a distance less than *ϵ* to a point in *X*, the Hausdorff distance between the two point clouds is less than *ϵ*.

7 If not, more work has to be done. See [4] for more details.

## References

1. J.B Ranck. “Head direction cells in the deep cell layer of dorsal presubiculum in freely moving rats”. In: Electrical Activity of Archicortex., G. Buzsáki, and Vanderwolf, C. H., eds. (1985), pp. 217–220.

2. M. Tsodyks, T. Kenet, A. Grinvald, and A. Arieli. “Linking spontaneous activity of single cortical neurons and the underlying functional architecture”. In: Science 286.5446 (Dec. 1999), pp. 1943–1946.

3. John P. Cunningham and Byron M. Yu. “Dimen-sionality reduction for large-scale neural recordings”. In: 17 (Aug. 2014). Review Article, 1500 EP -. url: http://dx.doi.org/10.1038/nn.3776.

4. Vin de Silva, Dmitriy Morozov, and Mikael Vejdemo-Johansson. “Persistent Cohomology and Circular Coordinates”. In: Discrete & Computational Geometry 45.4 (2011), pp. 737–759. issn: 1432-0444. doi: 10.1007/s00454-011-9344- x. url: http://dx.doi.org/10.1007/s00454-011-9344-x.

5. Mikael Vejdemo-Johansson, Florian T. Pokorny, Primoz Skraba, and Danica Kragic. “Cohomological learning of periodic motion”. In: Applicable Algebra in Engineering, Communication and Computing 26.1 (Mar. 2015), pp. 5–26. issn: 1432-0622. doi: 10.1007/s00200-015-0251-x. url: https://doi.org/10.1007/s00200-015-0251-x.

6. Gard Spreemann, Benjamin Dunn, Magnus Bakke Botnan, and Nils A. Baas. Using persistent homology to reveal hidden information in neural data. Version 1. 2015. arXiv: 1510.06629 [q-bio.NC]. url: https://arxiv.org/abs/1510.06629.

7. Adrien Peyrache, Marie M. Lacroix, Peter C. Petersen, and György Buzsáki. “Internally organized mechanisms of the head direction sense”. In: Nat Neurosci 18.4 (Apr. 2015). Article, pp. 569–575. issn: 1097-6256. url: http://dx.doi.org/10.1038/nn.3968.

8. Adrien Peyrache and György Buzsáki. Extracellular recordings from multi-site silicon probes in the anterior thalamus and subicular formation of freely moving mice. 2015. doi: 10.6080/K0G15XS1.

9. William E. Skaggs, Bruce L. McNaughton, and Katalin M. Gothard. “An Information-Theoretic Approach to Deciphering the Hippocampal Code”. In: Advances in Neural Information Processing Systems 5. Ed. by S. J. Hanson, J. D. Cowan, and C. L. Giles. Morgan-Kaufmann, 1993, pp. 1030–1037. url: http://papers.nips.cc/paper/671-an-information-theoretic-approach-to-deciphering-the-hippocampal-code.pdf.

10. P. McCullagh and J.A. Nelder. Generalized Linear Models, Second Edition. Chapman and Hall/CRC Monographs on Statistics and Applied Probability Series. Chapman & Hall, 1989. isbn: 9780412317606.

11. J. S. Taube, R. U. Muller, and J. B. Ranck. “Head-direction cells recorded from the post-subiculum in freely moving rats. II. Effects of environmental manipulations”. In: J. Neurosci. 10.2 (Feb. 1990), pp. 436–447.

12. Ryan M. Yoder, James R. Peck, and Jeffrey S. Taube. “Visual Landmark Information Gains Control of the Head Direction Signal at the Lateral Mammillary Nuclei”. In: Journal of Neuroscience 35.4 (2015), pp. 1354–1367. issn: 0270-6474. doi: 10.1523/JNEUROSCI.1418-14.2015. eprint: http://www.jneurosci.org/content/35/4/1354.full.pdf. xurl: http://www.jneurosci.org/content/35/4/1354.

13. S. J. Mizumori and J. D. Williams. “Directionally selective mnemonic properties of neurons in the lateral dorsal nucleus of the thalamus of rats”. In: J. Neurosci. 13.9 (Sept. 1993), pp. 4015–4028.

14. J. P. Goodridge et al. “Cue control and head direction cells”. In: Behav. Neurosci. 112.4 (Aug. 1998), pp. 749–761.

15. K Zhang. “Representation of spatial orientation by the intrinsic dynamics of the head-direction cell ensemble: a theory”. In: Journal of Neuroscience 16.6 (1996), pp. 2112–2126. issn: 0270-6474. eprint: http://www.jneurosci.org/content/16/6/2112.full.pdf. url: http://www.jneurosci.org/content/16/6/2112.

16. Yuri Dabaghian, Vicky L. Brandt, and Loren M. Frank. “Reconceiving the hippocampal map as a topological template”. In: eLife 3 (Aug. 2014). Ed. by Howard Eichenbaum. 25141375[pmid], e03476. issn: 2050-084X. doi: 10.7554/eLife.03476. url: http://www.ncbi.nlm.nih.gov/pmc/articles/PMC4161971/.

17. James J. Knierim, Hemant S. Kudrimoti, and Bruce L. McNaughton. “Interactions Between Idiothetic Cues and External Landmarks in the Control of Place Cells and Head Direction Cells”. In: Journal of Neurophysiology 80.1 (1998), pp. 425–446. issn: 0022-3077. eprint: http://jn.physiology.org/content/80/1/425.full.pdf. url: http://jn.physiology.org/content/80/1/425.

18. Adrien Peyrache, Natalie Schieferstein, and György Buzsáki. “Transformation of the head-direction signal into a spatial code”. In: bioRxiv (2017). doi: 10.1101/075986. eprint: https://www. biorxiv. org/content/early/2017/09/01/075986.full.pdf. url: https://www.biorxiv.org/content/early/2017/09/01/075986.

19. Mattia Rigotti et al. “The importance of mixed selectivity in complex cognitive tasks”. In: Nature 497.7451 (May 2013). Article, pp. 585–590. issn: 0028-0836. url: http://dx.doi.org/10.1038/nature12160.

20. Gurjeet Singh et al. “Topological analysis of population activity in visual cortex”. In: J Vis 8.8 (June 2008). 18831634[pmid], pp. 11.1–1118. issn: 1534-7362. doi: 10.1167/8.8.11. url: http://www.ncbi.nlm.nih.gov/pmc/articles/PMC2924880/.

21. Y. Dabaghian, F. Mémoli, L. Frank, and G. Carlsson. “A Topological Paradigm for Hippocampal Spatial Map Formation Using Persistent Homology”. In: PLOS Computational Biology 8.8 (Aug. 2012), pp. 1–14. doi: 10.1371/journal. pcbi. 1002581. url: https://doi.org/10.1371/journal.pcbi.1002581.

22. Carina Curto and Vladimir Itskov. “Cell Groups Reveal Structure of Stimulus Space”. In: PLOS Computational Biology 4.10 (Oct. 2008), pp. 1–13. doi: 10.1371/journal.pcbi.1000205. url: https://doi.org/10.1371/journal.pcbi.1000205.

23. Chad Giusti, Eva Pastalkova, Carina Curto, and Vladimir Itskov. “Clique topology reveals intrinsic geometric structure in neural correlations”. In: Proc Natl Acad Sci U S A 112.44 (Nov. 2015). 26487684[pmid], pp. 13455–13460. issn: 0027-8424. doi: 10. 1073/pnas. 1506407112. url: http://www.ncbi.nlm.nih.gov/pmc/articles/PMC4640785/.

24. Jose A. Perea. Multi-Scale Projective Coordinates via Persistent Cohomology of Sparse Filtrations. Version 2. 2016. arXiv: 1612.02861 [math.AT]. url: https://arxiv.org/abs/1612.02861.

25. A. Hatcher. Algebraic Topology. Algebraic Topology. Cambridge University Press, 2002. isbn: 9780521795401. url: https://www.math.cornell.edu/∼hatcher/AT/ATpage.html.

26. Robert Ghrist. “Barcodes: the persistent topology of data”. In: Bull. Amer. Math. Soc. (N.S.) 45.1 (2008), pp. 61–75. issn: 0273-0979. doi: 10. 1090/S0273-0979-07-01191-3. url: http://dx.doi.org/10.1090/S0273-0979-07-01191-3.

27. H. Edelsbrunner and J. Harer. Computational Topology: An Introduction. Applied Mathematics. American Mathematical Society, 2010. isbn: 9780821849255. url: http://www.ee.oulu.fi/research/imag/courses/Vaccarino/EdelsηBook.pdf.

28. Robert Ghrist. Elementary Applied Topology. CreateSpace, 2014. isbn: 1502880857. url: https://www.amazon.com/Elementary-Applied-Topology-Robert-Ghrist/dp/1502880857.

29. Gunnar Carlsson. “Topology and data”. In: Bulletin of the American Mathematical Society 46.2 (2009), pp. 255–308. url: https://doi.org/10.1090/S0273-0979-09-01249-X.

30. Pavan Ramkumar. pyglmnet. GitHub. 2016. url:https://github.com/glm-tools/pyglmnet.

31. Elmar de Koning. psimpl. 2011. url: http://psimpl.sourceforge.net/radial-distance.html.

32. Jennifer Kloke and Gunnar Carlsson. Topological De-Noising: Strengthening the Topological Signal. Version 2. 2010. arXiv: 0910.5947 [cs.CG]. url: https://arxiv.org/abs/0910.5947.

33. Jeffery Kline. LSQR. 2009. url: http://pages. cs.wisc.edu/∼kline/cvxopt.

34. Frédéric Chazal, Steve Y. Oudot, Marc Glisse, and Vin De Silva. The Structure and Stability of Persistence Modules. SpringerBriefs in Mathematics. nSpringer Verlag, 2016, pp. VII, 116. url: https://link.springer.com/book/10.1007/978-3-319-42545-0.

35. Ulrich Bauer. Ripser. GitHub. 2016. url: https://github.com/Ripser/ripser.

